# Software and pipelines for registration and analyses of rodent brain image data in reference atlas space

**DOI:** 10.1101/2025.04.25.650563

**Authors:** Maja A. Puchades, Sharon C. Yates, Gergely Csucs, Harry Carey, Arda Balkir, Trygve B. Leergaard, Jan G. Bjaalie

## Abstract

Advancements in methodologies for large-scale acquisition of high-resolution serial microscopy image data have opened new possibilities for experimental studies of cellular and subcellular features across whole brains in animal models. There is a high demand for open-source software and workflows for automated or semi-automated analysis of such data, facilitating anatomical, functional, and molecular mapping in healthy and diseased brains. These studies share a common need to precisely identify, visualize, and quantify the location of observations within anatomically defined regions, ensuring consistent and reproducible interpretation of anatomical locations and thereby allowing meaningful comparisons of results across multiple independent studies. Addressing this need, we have developed a suite of desktop and web-applications for registration of serial brain section images to three-dimensional brain reference atlases (QuickNII, VisuAlign, WebAlign, WebWarp, and DeepSlice) and for performing data analysis in a spatial context provided by an atlas (Nutil, QCAlign, SeriesZoom, LocaliZoom, and MeshView). The software can be utilized in various combinations, creating customized analytical pipelines suited to specific research needs. The web-applications are integrated in the EBRAINS research infrastructure and coupled to the EBRAINS data platform, establishing the foundation for an online analytical workbench. We here present our software ecosystem, exemplify its use by the research community, and discuss possible directions for future developments.

## Introduction

Animal models are invaluable resources for exploring brain architecture, mapping the distribution of cells types and molecules in the brain, and understanding the functional roles of brain structures. When combined with genetic or other modifications for mimicking human disease mechanisms, they enable the testing of hypotheses related to studies of health and disease. Widely accessible histological staining and microscopy techniques offer numerous benefits. They support spatial analysis since they preserve intrinsic brain architecture, and they are compatible with diverse labelling techniques to reveal specific cellular and molecular features. In tract tracing experiments, axonal tracers are injected into the brain, and transported through fibres, resulting in labelling of connections ^1^. Techniques like immunohistochemistry, immunofluorescence multiplexing, or in-situ hybridization reveal the distribution of specific cell types, gene expression patterns, and DNA sequences, for characterizing the brains of animal models.

Analysis of serial section images has traditionally involved labor-intensive manual approaches for defining anatomical regions, supplemented by quantification of features using various methods, including signal thresholding or stereological methods. Recent advancements have leveraged machine-learning based approaches for feature extraction as well as digital three-dimensional (3D) brain reference atlases. The use of 3D atlases supports a more standardized approach for defining anatomical regions, but requires accurate spatial registration between experimental section images and atlas volumes, which can be challenging to achieve (see, e.g., ^2,3^).

Many open-source scripts and software are now available for researchers conducting studies on mice and rats, tailored to different data modalities. Some address solely image registration to a reference brain atlas^4-6^, while others offer complete analytical pipelines^7-10^. Most are implemented in Python, Matlab, or as Fiji plugins, often requiring coding skills, which can be a barrier to a wider user base. Recent developments incorporate machine-learning to automate registration to atlases^11^, aiming to enhance efficiency and reduce biases in the registration process. However, to date, such approaches are most effective with structurally intact serial sections cut in standard planes. Deep learning models still struggle with distorted or damaged tissue sections, which commonly occur. Machine-learning methods are also increasingly employed for image segmentation, automating and standardizing feature extraction processes^12,13^. While most software is developed in-house to suit specific projects, there are increasing efforts to make software tools more accessible to the scientific community by providing user interfaces, flexible functionalities, and user documentation. Atlas-based software is thus increasingly employed in experimental studies, providing more efficient and standardized analyses compared to traditional methods (for review, see ^2,14^). Despite progress, it remains challenging to find combinations of user-friendly tools that can be tailored to the highly diverse needs of current research.

To facilitate standardized analyses using reference brain atlases, we here present our suite of interactive software and web-applications for atlas-based analyses of serial section images from rodent brains. The software has been designed to lower barriers for users, enabling image registration to atlases (QuickNII, VisuAlign, WebAlign, WebWarp, DeepSlice), diverse analyses in an atlas context (Nutil, QCAlign, LocaliZoom and MeshView), and inspection of sections and results in 2D and 3D reference space (SeriesZoom, LocaliZoom and MeshView). The tools are interoperable, comply with open and FAIR standards for research software ^15,16^, and shared through and integrated in the EBRAINS research infrastructure (https://ebrains.eu) with online user manuals and tutorials.

To demonstrate the capabilities of our software for inspecting, visualizing, and analysing brain section images from mouse and rat, we present three examples of use comprising datasets obtained from studies with unique experimental designs and varied analytical needs. In each example, we highlight the advantages of using the EBRAINS solutions while noting limitations and suggesting areas for future development. To our knowledge, this open-source software suite for serial section images is among the most comprehensive available. providing an adaptable analysis ecosystem for a wide range of study paradigms.

## Software overview

The software tools presented here are summarized in Table 1 with technical details and additional scripts provided in supplementary Table 1.

**Table 1.**
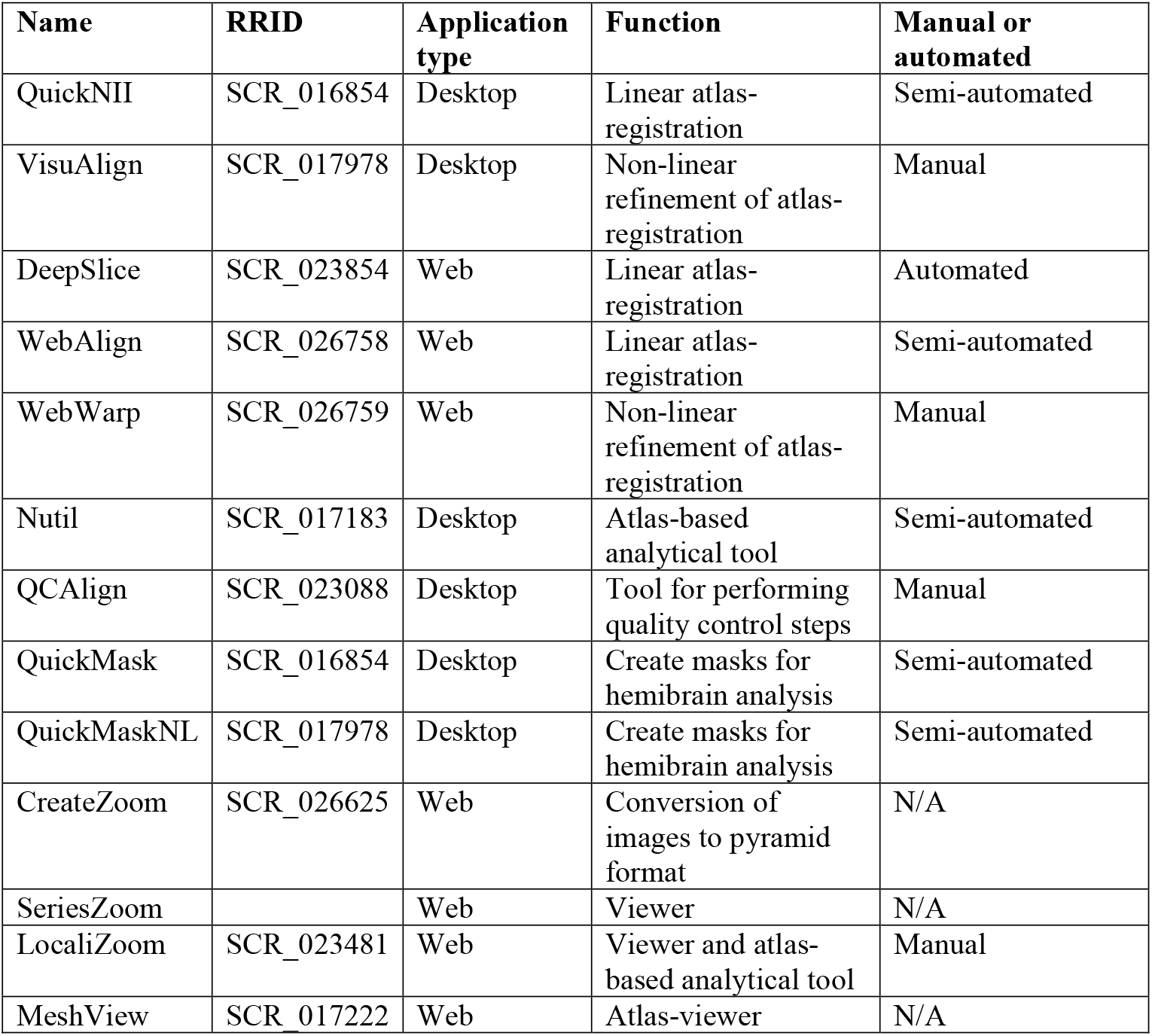
Overview of software.

### Software for image registration to a reference brain atlas

#### QuickNII

QuickNII is a standalone desktop application for user-guided affine spatial registration of brain section images to a 3D reference atlas. A key feature of the software is its ability to generate user-defined cut planes through the atlas templates that match the orientation of the cutting plane of the 2D experimental images, thereby generating adapted atlas maps. The reference atlas is transformed to match specific anatomical landmarks in the corresponding brain section images. In this way, the spatial relationship between the brain section image and the atlas is defined, without introducing transformations in the original image. Following registration of a limited number of sections containing key landmarks, transformations are propagated across the entire image series. As the propagation relies on the numbering of the section images, it is important that the sections are named using the file naming convention (_sXXX). The propagations should be validated and saved by the user for each section, with application of fine positional adjustments as required.

On the architecture level, QuickNII consists of two co-located executable components implemented in two programming languages for historical reasons. The GUI component is implemented in MXML+ActionScript (runs on Adobe Integrated Runtime, which is bundled as “captive runtime”, requiring no installation). A slicer service running in the background is implemented in Java (and has a bundled JRE requiring no installation). The two components communicate via standard output and local TCP connections (using the loopback interface).

#### VisuAlign

VisuAlign is a monolithic desktop application for applying user-guided nonlinear refinements (in-plane) to an existing, affine 2D-to-3D registration, such as one created using the QuickNII software. While linear registration tools are vital for bringing experimental image data to standardized coordinate spaces, for precise quantitative analysis residual anatomical variability among test subjects after registration must be addressed. VisuAlign is implemented in Java, with the graphical user interface built using JavaFX and FXML. While internally there are components and modules, like a slicer from QuickNII; the binary distributables are compiled with J-Link. This both eliminates the need for installing a separate Java Runtime and radically reduces the size of the VisuAlign package, at the same time rendering the internals inaccessible to the outside world. Thus, VisuAlign offers no other interfaces than the actual files that users load and save.

VisuAlign uses Delaunay triangulation over the target points (where the crosses have been dragged to), calculates barycentric coordinates for each target pixel inside their containing triangle, and then use the same coordinates in the source triangle (where the crosses have been dragged from) to sample the original (linear) atlas slice.

#### Deepslice

DeepSlice is a deep neural network, trained to predict the position of coronal mouse brain sections within the Allen mouse brain Common Coordinate Framework version 3 (CCFv3)^17^. DeepSlice produces registration files that are compatible with QuickNII and WebAlign. It performs a linear registration and does not predict non-linear deformations. DeepSlice is provided as both a Python package and as a web-application. This allows users to choose between a full featured version of the application and a simplified web user interface which prioritises ease of use.

#### WebAlign

WebAlign is a web-application for user-guided affine spatial registration of brain section image data (typically high-resolution histological images) to a 3D reference atlas. A key feature of the tool is its ability to generate user-defined cut planes through the atlas templates that match the orientation of the cutting plane of the 2D experimental images (atlas maps). Primarily it is a client-side web-application, running in a browser. Implementation languages are HTML5/CSS for the user interface, and JavaScript for processing. Requirements are deliberately kept low, WebAlign is expected to work properly with web browsers released in March 2017 or later [https://developer.mozilla.org/en-US/docs/Web/API/Fetch_API].

The server-side part of WebAlign consists of a very short PHP routine (10 instructions) implementing OIDC token exchange with the EBRAINS IAM service. This step is expected to remain necessary when integrating WebAlign into other environments, although the actual implementation language can be changed. Other requirements for the hosting server are minimal, at the time of writing, WebAlign requests whole files from its hosting server (compressed atlas packages are entirely transferred to the client).

Storage infrastructure of user data for WebAlign is integrated with the EBRAINS “Data-Proxy” (https://wiki.ebrains.eu/bin/view/Collabs/the-collaboratory/), that provides an S3-like interface. Adapting to similar infrastructures is expected to be straightforward. Feature requirements are again low since WebAlign can already operate with down/uploading whole objects at a time. WebAlign can use HTTP RANGE requests if users decide to store their DeepZoom images into ZIP archives (DeepZoom images may consist of several millions of small files, and they can be inconvenient to handle individually).

#### WebWarp

WebWarp is a web-application for nonlinear refinement of spatial registration of section images from rodent brains to reference 3D atlases. WebWarp is compatible with registration performed with WebAlign. Different experimental datasets registered to the same reference atlas allows spatially integration, analysis and navigation of these datasets within a standardised coordinate system. Technical details for WebWarp are identical to WebAlign: they operate on the same data and pose the same requirements towards both the browser and the backend (hosting server for the application and storage infrastructure for user data).

### Software for atlas-based analytical tools

#### Nutil

Nutil is a desktop application for pre- and post-processing of brain section images, which are typically large and difficult to process with standard image analysis software. It incorporates functionality for transforming, renaming and converting image file formats, and for quantifying features in the context of a reference brain atlas. It is a standalone Windows 64-bit application written in Qt C++.

#### QCAlign

QCAlign is a desktop application implemented in Java, the user interface is JavaFX and FXML. Internally it shares large amounts of code with VisuAlign: the two applications share their data descriptor format, the images and atlas overlays, which they load and display in a similar way (QCAlign is capable of working with both linear and nonlinear registrations). However, the actual functionality has a completely distinct implementation. New modules are implemented for enhanced atlas management (QCAlign allows users to customize atlas-granularity via a collapsible hierarchy-tree); for generating, pre-filling and interaction with a sampling grid; and for creating the statistical output.

#### QuickMask

QuickMask is a standalone software with a rudimentary GUI, enabling users to automatically generate customized hemisphere masks corresponding to brains sections registered to a reference atlas using QuickNII or VisuAlign. QuickMaskNL generates hemisphere masks which respect nonlinear deformations applied around the midline with VisuAlign. The customized masks are directly compatible with the Nutil software and can be used to perform separate analysis of the right and left hemisphere in brain sections. Preliminary versions of QuickMask and QuickMaskNL are available for download through the NITRC page for QuickNII and VisuAlign respectively.

### Viewers and related tools

#### CreateZoom

CreateZoom is a web-application for converting individual 2D images (.tiff; .jpeg or .png) to image pyramid files in Deep Zoom format (DZI): a prerequisite for the online QUINT service and for creating SeriesZoom and LocaliZoom viewers links. The application consists of a back-end service for concurrent batch creation of Deep Zoom Images via PyVips, a wrap of the lower-level image processing library libvips ^18^. The front-end allows users to define location and to select files to process. Internally, the application handles all actions asynchronously, to schedule processes and poll status of tasks, allowing the freedom to scale variables for the host. The storage service for such pyramid image files is connected from the EBRAINS “Data-Proxy” application accessed with a token acquired from the identity and access management service (IAM) at EBRAINS (https://wiki.ebrains.eu/bin/view/Collabs/the-collaboratory/). Processed images are compressed into .dzip files ready for further use.

#### SeriesZoom

SeriesZoom is the most basic online viewer we provide, allowing viewing of all DZI images (e.g. created by CreateZoom) from a single location. It does not show any atlas overlay or require any kind of series descriptor; it simply collects all DZI images it finds at a provided location. A filmstrip is provided with small icons, and the selected image is displayed in a pan-and-zoom fashion. SeriesZoom is implemented using HTML5, JavaScript, and CSS and runs entirely in the browser. It needs access to a series of DZI images. The current implementation assumes S3-like storage, with support for listing objects with a prefix. The discovery function can be easily adapted to different systems providing similar listing functionality. If there are no options for discovery, it can instead be modified to fetch an actual text/JSON with the list of DZI images.

#### LocaliZoom

LocaliZoom is a web-based pan-and-zoom 2D image viewer coupled with a volumetric atlas slicer, and a navigational aid showing the entire brain section image series as a “filmstrip”. Building on the open standard Deep Zoom Image (DZI) format, it efficiently visualises large brain section images in the gigapixel range, allowing image zoom from common, display-sized overview resolutions down to the microscopic resolution without downloading the underlying image dataset. In addition, LocaliZoom has an annotations and extraction feature. Markers can be manually placed by the user through the GUI, these markers are converted to point clouds easily visualised in the 3D viewer, MeshView and can be downloaded as csv files. LocaliZoom is also used as a brain section image viewer on the EBRAINS data sharing infrastructure for publicly shared dataset which have been registered to a reference atlas using QuickNII or VisuAlign.

LocaliZoom is a 100% client-side web application (HTML5, JavaScript, CSS); it is the base of WebWarp, without the need for a read-write storage system. Thus, its structure and requirements are similar to WebWarp. LocaliZoom needs images in DZI format, and a series descriptor that contains their relation to anatomical atlases. While LocaliZoom is simpler than WebWarp, it is also more flexible, a single “configuration.js” allows tuning all aspects of the application related to accessing data (series descriptor, DZI descriptor, DZI tiles), that already enabled us to use LocaliZoom with multiple image storage strategies (internal solution, swift object storage, EBRAINS “Data-Proxy”, individual tiles or tiles bundled into a single file), and descriptor formats (QuickNII, VisuAlign, WebAlign/WebWarp).

#### MeshView

MeshView is a web-application for real-time 3D display of surface mesh data representing structural parcellations from volumetric atlases, such as the Waxholm Space Atlas of the Sprague Dawley Rat Brain.

MeshView runs entirely in the web browser; and besides HTML5, JavaScript, and CSS, uses WebGL (a variant of OpenGL that is designed web environment, and programmable from JavaScript). MeshView is the most resource-hungry web application. One aspect is the network traffic it generates: while we provide highly optimized and compressed meshes, even the simplest atlas with 80 anatomical regions (WHS SD Rat version 2, from 2015)^19^ starts its operation by downloading 56 megabytes of data – other atlases are larger. The tool also supports loading point clouds (both as URL parameter and entered manually by the user), and the size of point cloud descriptors can also go up to tens or hundreds of megabytes.

The other resource-requiring aspect is rendering. Solid and transparent rendering of anatomical regions is fast (solid rendering uses simple Phong shading, and transparent rendering does not have to deal with complex order-independent algorithms as users find too many structures distracting and simply switch them off). Point cloud rendering is reasonably performant; users can adjust point size on a per-cloud basis. Lowering point size to a single pixel for clouds with huge amounts of points usually works well, but computers with integrated graphics solutions may experience slowdowns. The most complex feature is the support for cutting meshes as if they were solid objects. The algorithm is simple and elegant: the “even-odd rule” for filling closed polygons is extended to 3D (and thus filling polyhedra), and the cut plane (described by a point and a normal vector) is used to leave the surface open, effectively in “odd” state, stored in the stencil buffer. Almost every aspect of the algorithm is fast, but the stencil buffer must be restored to its initial, clear state between rendering each anatomical structure: we find that this single step is taxing for integrated graphics processors.

#### Additional scripts

Some additional scripts are available for QuickNII (https://github.com/Neural-Systems-at-UIO/QuickNII-extras) related to the propagation algorithm used during image registration to the reference atlas.

#### Python scripts

Python scripts for analysing point clouds related to the second example of use in the results part are available on Github (https://github.com/Neural-Systems-at-UIO/3d-point-clouds).

#### Overview of atlas versions currently available

Our applications have been extensively tested and support the following atlases: the Allen Mouse Brain Atlas CCF version 3, delineations from 2015 and 2017 ^17^; the Kim Unified Mouse Brain Atlas ^20^; DeMBA, a developmental mouse brain atlas for ages P4 – P56 ^21^ and the Waxholm Space Atlas of the Sprague Dawley rat v2 ^19^, v3 ^22^, v4 ^23^. In addition, some alternative atlases have been compiled in our tools (for details see https://quint-workflow.readthedocs.io/en/latest/QUINTintro.html#supported-atlases).

## Results

### The extended QUINT workflow

The QUINT workflow as originally presented in Yates et al.^3^ combined the use of three desktop applications supporting whole-brain mapping of labelled features in brain section image series from mice or rats. The core software in the workflow is QuickNII ^24^ for performing registration to a reference atlas, ilastik ^25^ for performing image segmentation for identifying features to be quantified, and Nutil for performing whole-brain mapping and regional feature quantification ^26^.The QUINT workflow has since been developed further with new releases of the core software offering new functionality and compatibility with alternative open-source software, and has been extended with new software.

We have added the possibility to use VisuAlign for refining the atlas registration by performing non-linear deformations of the atlas boundaries in the section image plane and QCAlign for performing quality control of the section image, to exclude brains regions from the analysis that correspond to damaged parts of the brain section and for defining the reference atlas hierarchical level to use for quantification, enabling greater flexibility ^27^. Optional functionality is provided using convolutional neural networks by DeepSlice, providing a fully automated registration of coronal mouse brain sections to the Allen CCFv3 atlas ^11^ compatible with the workflow.

For image segmentation and detection of the objects of interest, software such as QuPath ^28^, Cellpose ^29^ or Fiji ^30^ are now new options.

### Online web-applications

Following up on the requests and feedback from our users through the EBRAINS user support service, we have recently developed web-versions of the registration software, WebAlign (for linear registration) and WebWarp (for non-linear registration), available as Collaboratory apps on EBRAINS. The main advantages of the online applications are simplification of data management and file preprocessing steps which are a challenge when analysing large cohorts of mouse or rat brains. The file management is streamlined, enabling image series uploads via the Data Proxy service and conversion of the image files to pyramid files by the CreateZoom app. Taking advantage of the EBRAINS core services for user identification (IAM) and the Collaboratory service (https://wiki.ebrains.eu/bin/view/Main/), users get access to a private working space, where they can upload their datasets to a private bucket and use entire analytical pipelines, where image registration to a reference atlas of mouse or rat brain is the first step. We have provided a public demonstration collab to guide users (https://wiki.ebrains.eu/bin/view/Collabs/image-registration-and-analysis-demo). After a brief introduction on how to setup all the web-applications and convert the image files to pyramid files (DZI), the online registration is done interactively in the web browser. All the results are saved in the Collaboratory storage bucket and can be opened in the WebWarp application for refinement with non-linear registration. WebWarp also uses the DZI file format, which allows users to zoom-in to high-resolution, to better decipher region boundaries and achieve a more precise registration. Furthermore, EBRAINS users can open datasets that have been publicly shared through EBRAINS by fetching a link on the dataset card. By opening this link in WebAlign, the image series is automatically created and saved as a WebAlign file (.waln) in the storage bucket. Users can then modify the registration or apply non-linear corrections using WebWarp. All the web-apps are designed to be interoperable and compatible with other analysis tools. They allow users to compare different datasets and combine the results from multimodal experimental setups, thus increasing the reusability of data.

The integrated app LocaliZoom enables visualization, annotation of registered image series and extraction of coordinate points which can be visualized in the MeshView app together with atlas meshes. In addition, high-resolution microscopy viewers such as SeriesZoom and LocaliZoom provide a seamless user experience and can generate image service links for datasets shared via the EBRAINS data sharing service, upon request by the data provider.

### Examples analyses

To date, our software has been cited in over 130 original research articles, ranging from tract tracing experiments to classical histological studies, and to tissue clearing experiments captured by light sheet microscopy. To illustrate the scope of the software for inspecting and analysing brain sections from mouse and rat, we present three examples comprising datasets obtained from studies with unique experimental design, requiring different solutions and analytical approaches. We highlight the advantages of using the EBRAINS solutions in each case; and point to limitations and areas for future development.

#### 1. Mapping of efferent connections in the rat brain

In a recent article, Reiten et al. ^31^ shared a large data collection from tract tracing experiments in rats aimed at investigating the connections between brain regions involved in spatial navigation, decision making and working memory. The collection consisted of serial coronal sections from 49 rat brains in which the anterograde tracers Biotinylated dextran amine (BDA) or *Phaseolus vulgaris*-leucoagglutinin (Pha-L) were injected in the orbitofrontal, parietal or insular cortex ^32-36^. Because publicly available rat connectivity datasets are scarce, the authors prepared the data and metadata for sharing through the EBRAINS data portal, with the collections shared as three datasets: projections from the insular cortex (https://doi.org/10.25493/WK4W-ZCQ); projections from the posterior parietal cortex (https://doi.org/10.25493/FKM4-ZCC); and projections from the orbitofrontal cortex (https://doi.org/10.25493/2MX9-3XF).

To increase interoperability and opportunities for combined analysis and reuse of the datasets, thereby improving FAIRness ^15^, all section images were registered to the 3D Waxholm Sprague Dawley rat atlas version 4 ^23^ using the registration software QuickNII ^24^ and VisuAlign. <u>The</u> workflow is presented in Figure 1a. First, preprocessing steps were applied to ensure that the section images had a proper orientation and sequential positioning along the antero-posterior axis by at least two researchers. Then, linear registration of the images to the 3D atlas was performed in QuickNII for a global positioning (Fig. 1b), before refinement of the registration using in-plane non-linear deformations with VisuAlign (Fig. 1c). Visual landmarks in the tissue are used to guide these deformations (Fig 1ii and 1iii). As illustrated in Figure 1, the non-linear adjustment of the atlas to match the experimental sections was essential to achieve correct positioning of the labelled fibres, especially for smaller brain regions or specific nuclei, like the submedius thalamic nucleus (Fig 1i and 1iv). The registration software finally allowed users to create atlas overlays on their images, downloadable as 2D images. These registration processes still rely on manual work and good knowledge of the brain anatomy by the users. We believe that development of automatic registration algorithms for the rat brain series will bring a welcome addition to this workflow.

**Figure 1.**
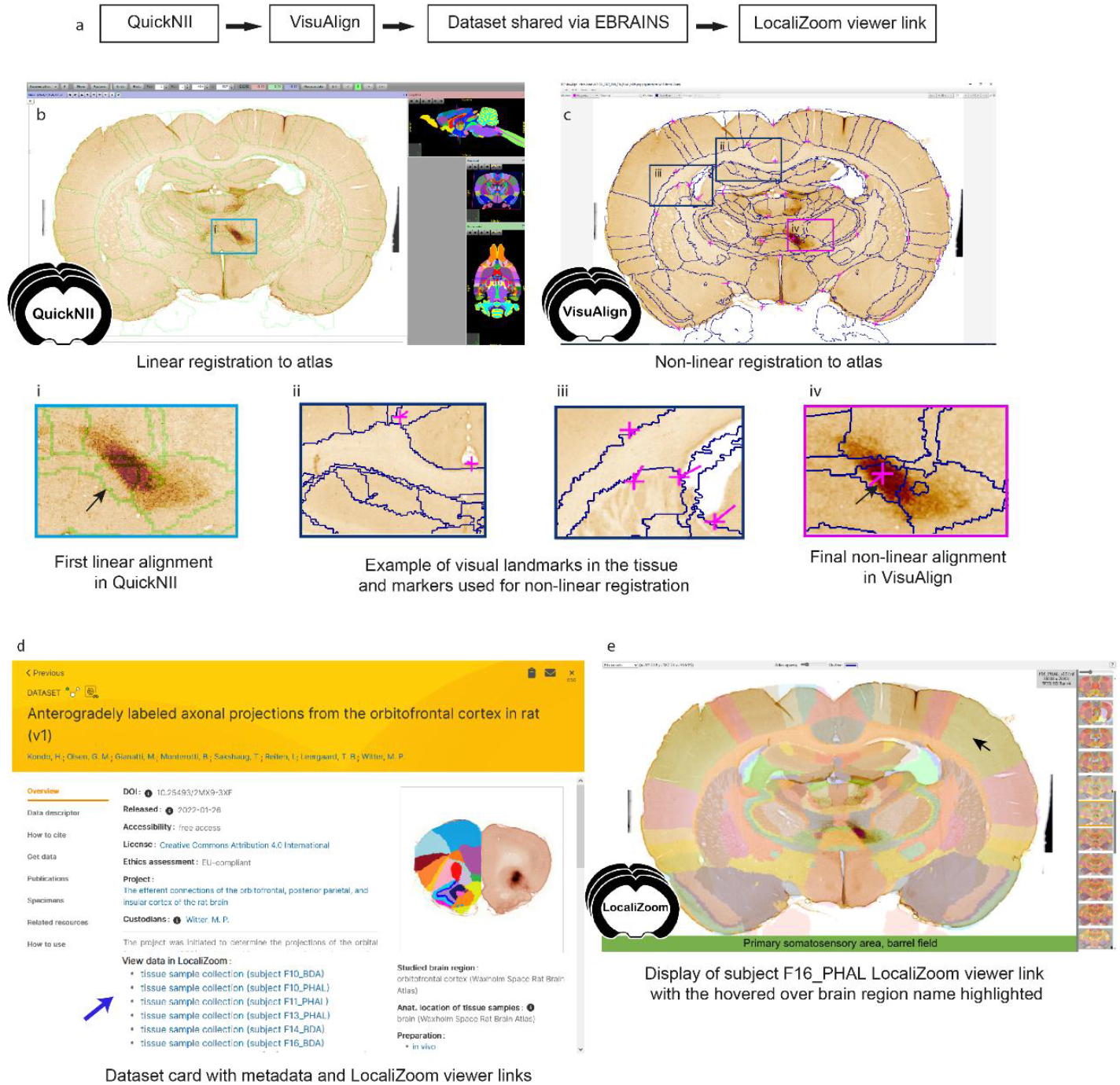
**(a) Workflow for section image registration to a reference atlas and creation of LocaliZoom viewer links. (b)** Section image registration with the QuickNII software and **(c)** VisuAlign software. **(ii and iii)** Tissue landmarks are used to guide the non-linear registration. Comparison of registration before **(i)** and after **(iv)** non-linear adjustments illustrates the importance of this step for correct alignment of small brain regions. **(d)** In this case, the atlas registrations were made available on the EBRAINS dataset card via the service link provided through the LocaliZoom app. **(e)** LocaliZoom service link displaying the whole brain section image series with the atlas overlay visible and brain region name is displayed when hovering the mouse on it for ease of identification. The data used here for illustration is from subject F16_PHAL and dataset 10.25493/2MX9-3XF.

For datasets published on EBRAINS (Fig 1d), users have the option to request viewer links on the EBRAINS data cards for displaying the section images with atlas overlays (Fig. 1e). The viewer links are created using our image viewer LocaliZoom, and allow users to 1) browse through the whole image collection arranged in anterio-posterior sequence; 2) zoom in on particular regions in the experimental images for closer inspection as they are stored in the pyramid file format DZI; 3) explore the images with or without the atlas overlays visible at different intensities 4) share the viewer links with their collaborators and colleagues.

#### 2. Mapping projections in the mouse brain

In this example, the authors were interested in the topographical organisation of the first link in the neuronal projections from the cerebral cortex to the cerebellum: the corticopontine projections ^37^. To study these connections in mice, they used publicly available serial brain section images, which had connections revealed using injection of fluorescently labelled anterograde tracer molecules. Images from the Allen Mouse Brain Connectivity collection and datasets published on EBRAINS were combined (https://doi.org/10.25493/GDYP-B1B); (https://doi.org/10.25493/11HT-S4B).

The workflow is illustrated in Figure 2a. To extract information about the location of these projections, all the brain section images were registered to the Allen mouse brain Common Coordinate Framework (CCFv3_2017) using the QuickNII software. The location of the axonal terminal fields in the pontine nuclei was recorded manually using a local instance of LocaliZoom (RRID:SCR_023481; https://localizoom.readthedocs.io) for assigning point coordinates at regular intervals (Figure 2b). This feature in LocaliZoom allows users to place markers on the objects-of-interest and thereby capture their coordinates (fig.2b’). The plotting reflects the observed density of labelling. Users can easily navigate through their image collections using the filmstrip view (Fig. 2b). A manual process for extracting coordinates was shown to be more precise in this case than a more automatic process using image segmentations (e.g. the QUINT workflow) as too much non-specific labelling was included (for details see ^37^).

**Figure 2.**
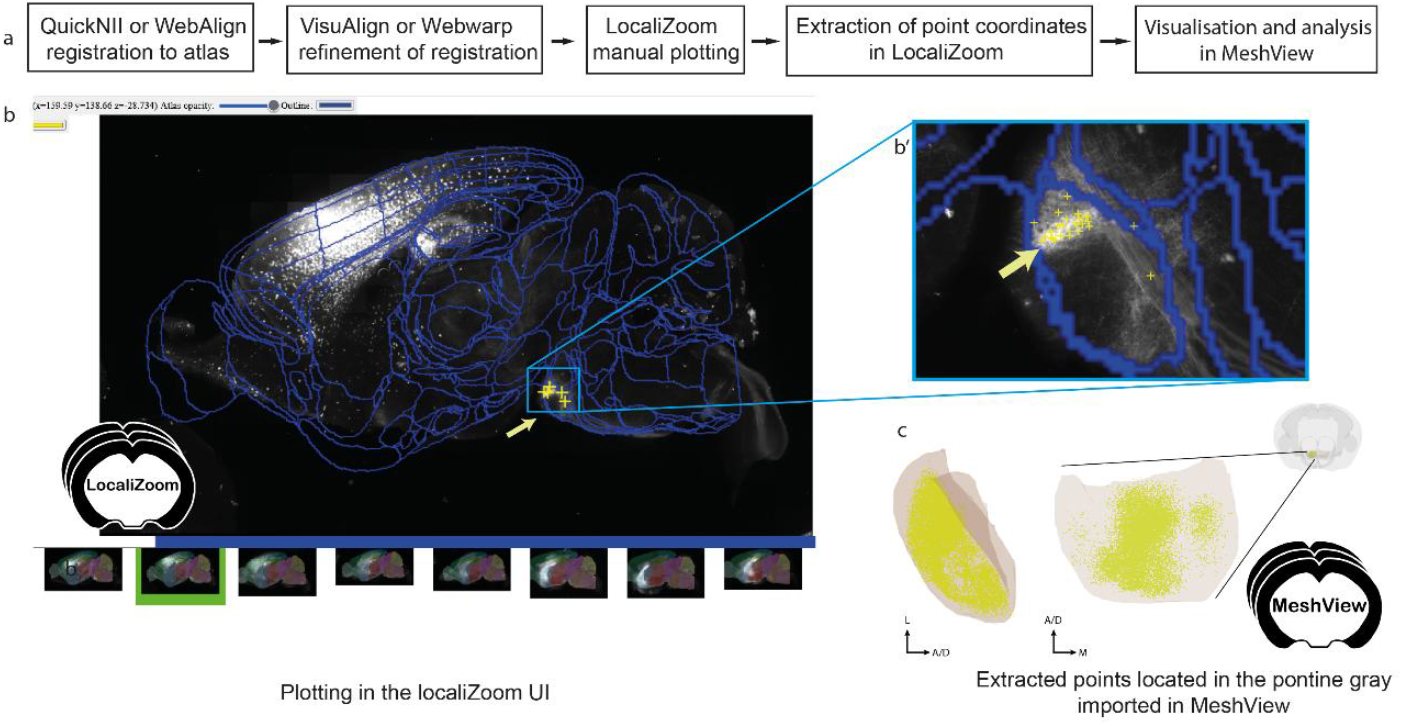
**(a) Workflow for extracting point clouds from registered brain section series. (b)** The LocaliZoom app displays the original experimental images after they have been converted to a pyramid format (DZI). **(b’)** Users can zoom in and out of their high resolution images. The atlas-registrations are imported from a JSON file generated by the registration software (either QuickNII/VisuAlign or WebAlign/WebWarp) and can be visualised as contour lines or coloured brain regions with different levels of transparency using a transparency slider button. The colour of the outlines can be changed by the user. **(c)** Extracted points can be visualised in MeshView. The data used here for illustration is from subject littermate control 11643_13 and dataset 10.25493/11HT-S4B.

The markers representing the labelling, generated in LocaliZoom, were converted to point clouds, and the output file from LocaliZoom was directly uploaded to the 3D viewer MeshView (RRID:SCR_017222; details; https://meshviewfor-brain-atlases.readthedocs.io/en/latest/) (Fig 2c). Some special features were added to MeshView to aid visualisation of patterns in the terminal distributions located in the pontine grey region across several animals (Figure 3).

**Figure 3.**
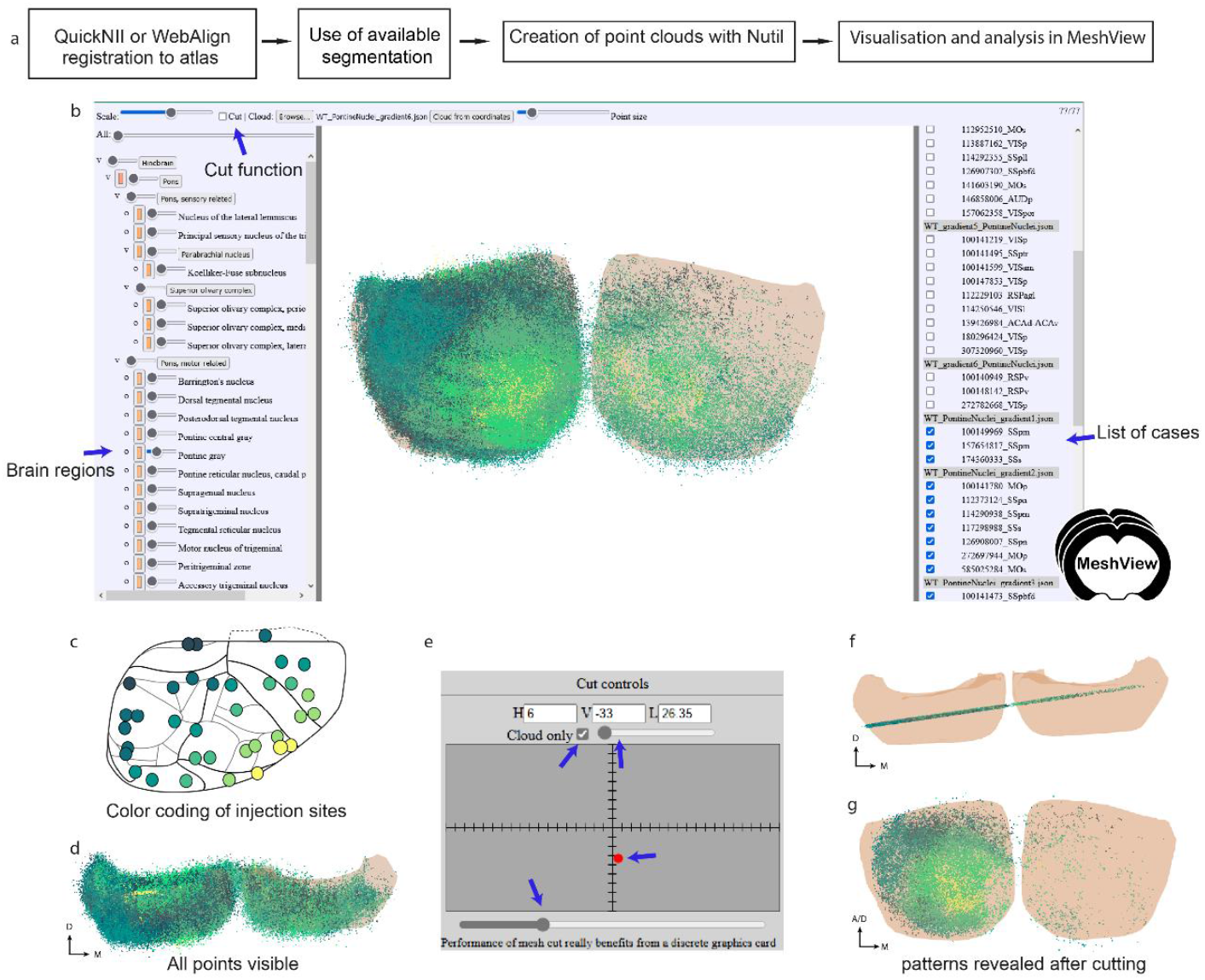
**(a) Workflow for extraction of point clouds from brain section series and analysis of topographical patterns in MeshView. (b)** The main UI of MeshView is shown in with selection of the brain region mesh on the left side (arrow pointing to the pontine grey region). The meshes have different degrees of transparency and are color-coded according to the atlas hierarchy label. These colors are editable by the user. The list of uploaded point cloud files (JSON format) is shown on the right side and can be interactively toggled on and off. **(d)** same point cloud as in **(b)** viewed from above illustrates the density of points. **(e)** panel showing the “double cut” feature implemented in MeshView for this specific study. It enables the creation of point cloud slices for an easier inspection of the topographical patterns **(f and g)**. The data used here for illustration is from point clouds showing spatial distribution of corticopontine projections originating from 35 tract-tracing experiments in wild type mouse (https://doi.org/10.25493/GDYP-B1B).

The cutting feature is found on the top left. **(c)** A schematic of the cortical tracer injection sites is shown where the positions of the injections have been color-coded from yellow to dark green based on the topographical location (see github repo https://github.com/Neural-Systems-at-UIO/3d-point-clouds for more details).

Indeed, when all point clouds from many animals are co-visualised (Fig 3b and d), no obvious pattern could be deciphered. By colour-coding the obtained point clouds according to the position of the injection site in the cortex, topographical patterns became visible (Fig 3c).

Jupyter notebooks with Python scripts were shared with the datasets allowing users to reproduce these patterns or apply the method to their own data (see the Methods part for more details). Additionally, a double cut feature was added to MeshView to visualise the point clouds as a slice (Fig 3e, f); thereby several different distributions patterns could be demonstrated like fan-like or concentric distributions (Figure g) (See Ovsthus et al. 2024 ^37^ for more details).

This type of result can be obtained by using the web applications: WebAlign, WebWarp and LocaliZoom in the EBRAINS Collaboratory (See https://wiki.ebrains.eu/bin/view/Collabs/image-registration-and-analysis-demo).

To our knowledge, no similar workflow is available today. Although it is a manual process, the main advantage is the possibility to extract signal where automatic image segmentation algorithms fail, i.e. when the signal to noise ratio is too low or when objects are too densely packed like the cell somas in the hippocampal principal layers.

#### 3. Characterising mouse brain composition in a complex high-throughput study

In a recent study, Gurdon et al. ^27^ investigated the effect of genetic makeup on the development of Alzheimer’s disease using a novel mouse model (AD-BXD), with a goal to reveal resilience mechanisms. The mouse model incorporated genetic diversity with AD risk mutations: resulting in strains with variable symptomatology despite carrying identical high-risk mutations ^38^. In an exploratory study of multiple AD-BXD strains, brains from 40 mice were divided into hemibrains: with one hemibrain dissected to supply tissue for bulk-RNA sequencing (Figure 4a). The other hemibrain was sectioned, labelled and imaged to allow histological brain-wide mapping of neurons, microglia, astrocytes and beta-amyloid (Figure 4b) (https://doi.org/10.25493/SZ0M-EE6). Due to the exploratory nature of the histological study, there was a need for a comprehensive analysis method supporting regional comparison across ages and strains. The use of a reference atlas to define regions for analysis supports such an exploratory approach. However, accurate registration is critical, and can be difficult to achieve and verify, especially for high-throughput studies involving genetically diverse animals as the anatomy of experimental models may differ from reference animals. Brain sections may also be distorted, damaged or torn by the sectioning process.

**Figure 4.**
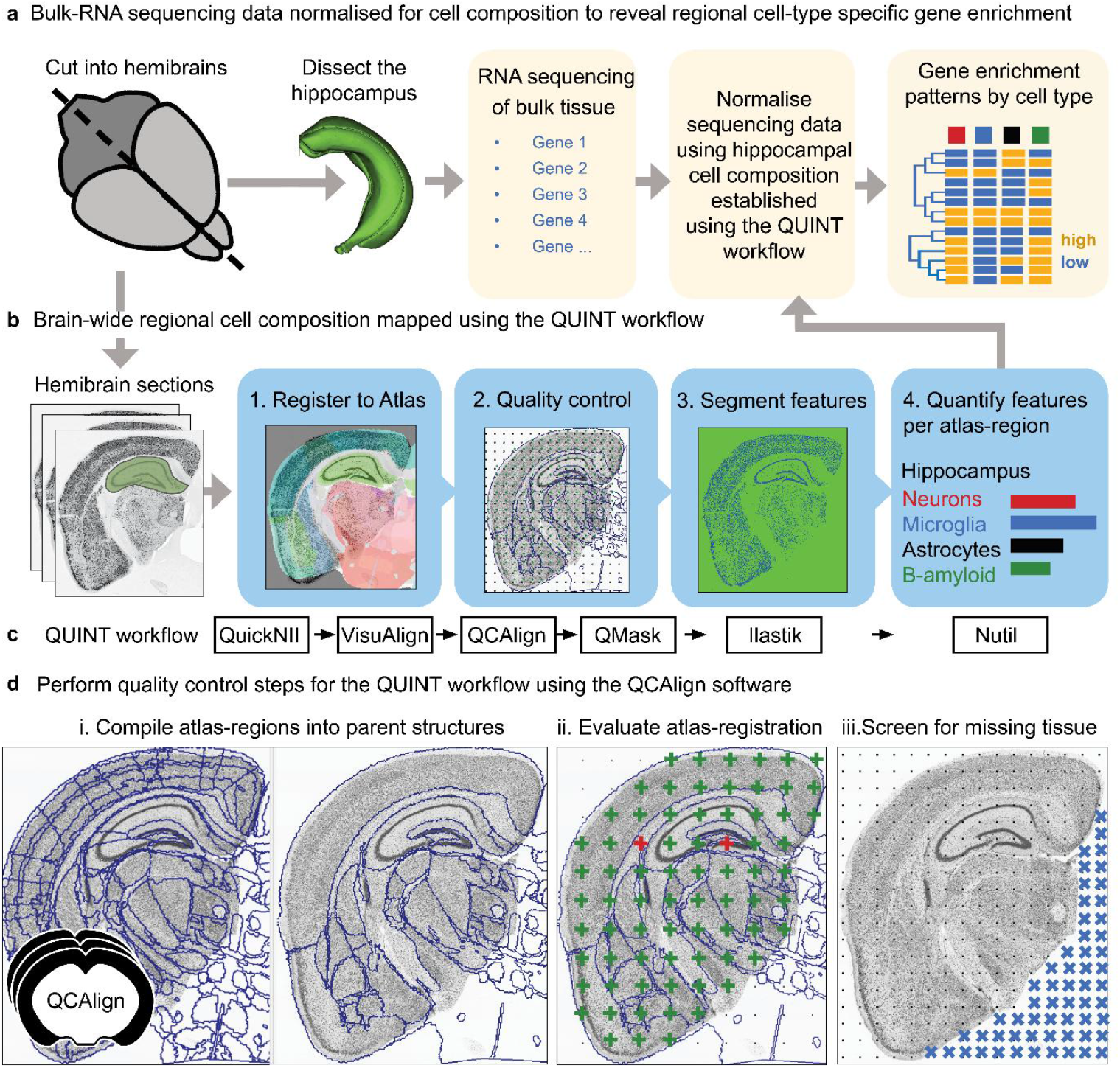
Forty AD-BXD mice were used to explore resilience mechanisms in Alzheimer’s disease using transcriptomic and histological methods. **(a)** To reveal cell-type specific gene set enrichment patterns in the hippocampus, brains from the forty mice were divided into hemibrains, with one hemibrain microdissected, supplying tissue for bulk-RNA sequencing. The bulk-RNA sequencing data was normalized using cell composition data from the hippocampus from the other hemisphere, established using the QUINT workflow. By integrating the bulk-RNA sequencing data with histological results from the other hemisphere, candidate genes and pathways involved in resilience to disease could be revealed in a region-specific as well as cell-type specific manner **(b and c)**. To explore cell composition using a reference brain atlas, histological data from one hemisphere was analyzed using the extended QUINT workflow **(c)**, quantifying neurons, microglia, astrocytes and beta-amyloid. This involved four key steps: 1. Registration to a reference brain atlas using QuickNII and VisuAlign. 2. Quality control steps using QCAlign with creation of hemibrain masks using QuickMask. 3. Image segmentation using ilastik to identify the features-of-interest. 4. Quantification of the features-of-interest in reference atlas regions using Nutil. **(d)** To ensure reliable histological results, the quality control software, QCAlign, was developed and integrated in the QUINT workflow providing functionality for i) establishing parent regions for enabling quality control of the atlas-registration and to use for quantification of features, ii) for evaluating the quality of the atlas-registration using systematic sampling (green crosses indicate correct registration, red crosses indicate incorrect registration), and iii) for establishing regions and sections to exclude from analysis due to damage (missing tissue represented by blue crosses).

In collaboration, we established the BRAINSPACE project for making the QUINT workflow applicable for such high-throughput applications ^27^. The QUINT workflow provided the functionality for bringing features into a common reference space: with registration to the Allen CCFv3 performed with QuickNII, with VisuAlign used to perform the manual adjustments needed to achieve a good anatomical fit over the sections, matching the outer edges as well as internal boundaries made visible by labelling (Figure 4c).

A limitation was a need to validate the region boundaries to be used for quantification due to the expected anatomical differences between individual mice. This validation was challenging, since not all regions included in the reference atlas could be identified in the section images of this study. To bypass this, reference regions of the atlas were combined into parent regions with visible boundaries in the section images. These parent regions were in turn used as boundaries for quantification. We developed the QCAlign software for compiling reference regions into suitable parent regions (Figure 4d, i), and for comparing these regions to boundaries visible in the section images by systematic sampling (Figure 4d, ii). By utilising QCAlign, we were able to confirm accurate registration of 77 parent regions across the whole brain, which were then used for quantification of labelled features. We also used QCAlign for rapidly screening sections for damage, allowing systematic removal of sections and regions with more than a set damage percentage (Figure 4d, iii).

By combining the use of VisuAlign for refining the atlas-registration with QCAlign for establishing regions with visible boundaries to use for quantification, for validating the atlas-registration to these regions, and for performing checks for tissue damage; we were able to counter the distortion and damage that is common in histological studies, ensuring reliable results. As the coronal sections originated from one hemisphere only, the QuickMask software supplied customized hemibrain masks tailored to each section and required by Nutil for hemibrain analysis.

By establishing regional cell composition in this study, results could be compared across ages and strains. Furthermore, the histological results could be used to normalize bulk-RNA sequencing data from corresponding regions from the other hemisphere, allowing candidate genes involved in resilience and progression of AD to be revealed in a region-specific as well as cell-type specific manner (Figure 4b). Thus, by integrating transcriptomic data with regional cell composition data, resilience pathways could be revealed and localized to specific cell-types. While this was a pilot study, focusing on the hippocampus for the development of the integration method, it demonstrates the power of integrating different data modalities using location as a common denominator.

## Discussion

To date, the software summarised here has been used in a wide range of published studies, using a range of animal models and experimental methods, and having a variety of output requirements. These studies have in common that they are based on 2D brain sections and aim to map the section images into a 3D common reference space. This integration facilitates data comparisons and lays the foundation for future discoveries. It also supports a more nuanced understanding of the brain and its diseases ^39^. We have showcased three examples of use of the software to meet challenges unique to different experimental setups. In the first, the desktop applications, QuickNII and VisuAlign are used for performing atlas-registration, enabling creation of LocaliZoom service links for displaying rat tract-tracing image series in the Waxholm Space atlas through the EBRAINS data sharing platform. The second demonstrates mapping of cortico-pontine connections in the mouse, with extraction of point coordinates of the terminal fields using the LocaliZoom application and study of topographical patterns using the MeshView software. Finally, the third demonstrates comprehensive brain-wide mapping of mouse brain data in a high-throughput context using the extended QUINT workflow, allowing location-based integration of data of two different modalities (histological and transcriptomic data).

Our applications have been widely used by the research community, with the software in the QUINT workflow used to provide brain-wide counts of various cell types, receptors and pathological markers in brain sections from traditional histological studies, ^40-49^; to investigate brain connectivity in tract-tracing experiments ^49-54^; and to map markers in tissue clearing experiments captured by light sheet microscopy^55,56^. While our software was not developed specifically for light sheet microscopy data, a 2D to 3D registration method has some advantages over 3D to 3D methods for this data type. This is because clearing procedures induce deformations in the tissue, which can be challenging to adjust for using volumetric registration approaches, ultimately affecting the subsequent registration result. Automated registration methods can be tailored to specific tissue clearing techniques^57^; however, the ability to interactively refine the atlas-registration using visual landmarks, as implemented in VisuAlign, is a clear advantage as it can be easily adopted without coding ability ^55^. Furthermore, the QUINT workflow provides a means for transparent analysis as the atlas-registration can be shared with the data and used with our web-applications (SeriesZoom and LocaliZoom) to create shareable microscopy viewer links with atlas overlays, as demonstrated for several datasets shared through the EBRAINS data sharing platform (ebrains.eu).

While most published studies have used our software as described in our user documentation, innovation is at play, with a proportion developing their own scripts and methods used in concert with the EBRAINS software to solve problems unique to their own experimental design. For example, the Henderson group have combined the use of QuickNII, VisuAlign, QuickMask and Nutil with the popular histopathology toolbox QuPath ^28^ for identifying features-of-interest, creating a modified version of the QUINT workflow for fluorescence ^40,41,58^. They have openly shared this method as an iprotocol, (doi: dx.doi.org/10.17504/protocols.io.4r3l22y6jl1y/v1). By sharing these scripts and methods, the research community benefits from access. Obstacles to analysis are also revealed, which can prompt developers to implement new features in their software, continuing the development cycle. Software development is thereby a community effort, with no clear start and end point. However, it is clear that by developing analytical tools in concert with a coordinated research infrastructure, as we have done here through the Human Brain Project with a continuation in the EBRAINS research infrastructure, we have been able to develop more mature research software, with more community involvement, than would have been possible without the research infrastructure approach. We have been able to offer web-applications through a research infrastructure for user authentication, resource allocation and data storage. We have also reached a larger target audience and have been able to offer a better user support service, which has provided the feedback needed to drive developments.

Currently, our applications incorporate two adult mouse brain atlases (Allen CCFv3^17^ and Kim Unified Mouse Brain Atlas^20^), one adult rat brain atlas (WHSSD)^23^, and a developmental mouse brain atlas for ages P4 – P56 (DeMBA)^21^. Brain atlas development is a growing field: with new atlases being released and existing atlases extended on a continuous basis. To assist developments using these resources, the BrainGlobe initiative provides an overview of available atlases for small animal models and have created an Atlas API which compiles these atlases and their metadata as a resource for developers (https://brainglobe.info/)^59^. We are in communication with them regarding future API developments, with potential to expand our atlas repertoire to match this collection in future releases. Co-development efforts such as these are critical to ensure compatibility of software across research groups and institutions; and to develop and adopt the standards needed to maximise the impact of such developmental efforts.

In a future development, we plan to release the entire QUINT workflow for brain-wide mapping in an online workbench environment. This will provide a streamlined user experience and ameliorate issues relating to file preparation and transfer between software, which can complicate offline workflows. It will also allow users to create shareable microscopy viewer links, increasing the FAIRness ^15^ of the datasets and related analyses. While online solutions have clear advantages over downloadable software, they also have downsides as they require on-going maintenance and are more prone to issues due to multiple dependencies. By providing both online and offline solutions using standardized atlases that are interoperable with solutions for transparent data sharing, our software promote data FAIRness and facilitate scientific discovery and data reuse.

## Acknowledgements

We thank our colleagues at the Neural Systems laboratory for their support, including Heidi Kleven, Ingvild Bjerke, Ingrid Reiten, Ulrike Schlegel, Camilla Blixhavn Hagen, Eszter A. Papp and Martin Øvsthus as well as all our fellow researchers and software users for their feature requests and feedback.

This work received funding from the European Union’s Horizon 2020 Framework Program for Research and Innovation under the Specific Grant Agreement No. 785907 (Human Brain Project SGA2), Specific Grant Agreement No. 945539 (Human Brain Project SGA3), the European Union’s Horizon Europe Programme for Research Infrastructures under the Specific Grant Agreement No. 101147319 (EBRAINS 2.0), and The Research Council of Norway under Grant Agreement No. 269774 (INCF Norwegian Node).

